# Pathological LSD1 mutations repress enhancer-mediated gene regulatory networks in early differentiation

**DOI:** 10.1101/2021.08.11.455900

**Authors:** Daria Bunina, Pierre-Luc Germain, Alejandro Lopez Tobon, Nadine Fernandez-Novel Marx, Thomas Dahlet, Christian Arnold, Andrew Ahern, Annique Claringbould, Mang Ching Lai, Sampath Rangasamy, Vinodh Narayanan, Peter Lorentzen, Karen M. Park, Antonio Adamo, Giuseppe Testa, Judith Zaugg, Kyung-Min Noh

**Affiliations:** European Molecular Biology Laboratory (EMBL), Genome Biology Unit, Heidelberg, Germany; European Molecular Biology Laboratory (EMBL), Structural and Computational Biology Unit, Heidelberg, Germany; Department of Experimental Oncology, European Institute of Oncology, IRCCS, Milan, Italy; Center for Rare Childhood Disorders and Neurogenomics Division Translational Genomics Research Institute, Phoenix, USA; Department of Economics, University of San Francisco, San Francisco, California, USA; Citizen scientist, San Francisco, California, USA

## Abstract

Lysine-specific demethylase 1 (LSD1/KDM1A) demethylates both histone and non-histone substrates, recruits repressive chromatin complexes, and is increased in cancers. De novo LSD1 mutations impairing protein function lead to a rare developmental disorder, but the molecular details of the pathology remains unclear. Using patient-derived fibroblasts, reprogrammed pluripotent stem cells, and differentiated cells, we found over 4000 differentially expressed genes and 68 transcription factors (TFs) whose motif accessibilities changed upon LSD1 mutation. An enhancer-mediated gene regulatory network approach identified transcriptional repressors with impaired activity in fibroblast and stem cells, leading to erroneous activation of their target genes. We also revealed overall decreases in TF target gene expression during early lineage differentiation of LSD1 mutant stem cells, likely caused by increased activity of repressive histone deacetylases (HDACs), co-factors of LSD1. Indeed, an HDACs inhibitor restored changes in gene expression including downregulation phenotype. Our findings characterize the molecular pathogenesis of LSD1 mutations and suggest potential therapeutic strategies for the developmental disorder and cancers caused by LSD1 dysregulations.

## Introduction

Epigenetic modifications on chromatin are essential for regulating cell type specific gene expression and, therefore, normal development. The positioning and dynamics of histone lysine methylation at enhancers and promoters, carried out by lysine methyltransferases and demethylases, is critical for cellular processes, such as transcription^1^, DNA repair^2^, the cell cycle^3^, and differentiation capacity^4^. Lysine-specific demethylase 1 (LSD1, encoded by the KDM1A gene) demethylates histone H3 lysine 4 mono- and di-methylated moieties (H3K4me1/2)^5^, marks found at active and poised enhancers and promoters. LSD1 also recruits repressive protein complexes, such as CoREST^6^ and NuRD^7^, to chromatin. LSD1 has been implicated in the control of cell differentiation^8^ and early embryogenesis^9^ possibly by silencing active enhancers in stem cells when they undergo differentiation. Functional alterations of LSD1 have been found in pathological conditions, such as hematopoiesis impairment^10^ and various cancers^11^.

Dominant missense point mutations in LSD1 cause the developmental disorder^12^ Cleft Palate, Psychomotor Retardation and Distinctive Facial Features (CPRF), which resembles Kabuki syndrome, characterized by distinctive facial features, muscle and skeletal abnormalities, and cognitive impairment. Three of the pathological mutations (Glu403Lys, Asp580Gly, Tyr785His) in purified proteins have been shown to impair LSD1 catalytic activity and, to varying extents, LSD1 binding to the histone H3 N-terminal tail^13^. LSD1 inactive mutants were also able to recruit corepressors (e.g., CoREST, HDAC1/2) when stably expressed in transfected cells^13^. Nevertheless, the molecular mechanisms by which the LSD1 mutations cause pathological manifestation have not been elucidated in the context of patient cells.

Here, using patient material, we investigated the effects of the pathological mutations Glu403Lys and Tyr785His on chromatin and gene expression in different cell types and across differentiation. We observed widespread misregulation of gene regulatory networks and increased transcriptional repression during differentiation, likely caused by increased activity of LSD1 co-factors, such as HDACs. Treating the cells with an inhibitor of HDACs, an FDA approved drug, rescued the molecular phenotype of the cells. Even with a limited number of patient cases, our integrated data analysis has identified a target pathway that may help treat a rare human disease. Our findings also provide a genome regulatory feature coupled to LSD1 dysregulations that may apply to various cancers.

## Results

### LSD1^MUT^ impairs chromatin accessibility at regulatory elements

To understand how de novo heterozygous mutations in LSD1 cause CPRF syndrome, we obtained primary skin fibroblasts, which express LSD1 (Extended Data Fig.1a), from two male pediatric patients (family A - Glu403Lys and family B - Tyr785His), and their healthy fathers (Fig. 1a). Both mutations maintain the overall amino acid backbone structure, but change polar groups to basic. Given LSD1 largely acts on chromatin at regulatory elements, such as enhancers and promoters, we hypothesized that heterozygous mutant cells would exhibit a disruption of chromatin at regulatory elements. To test this idea, we used ATAC-seq to map chromatin accessibility at regulatory elements in both the LSD1 mutant (LSD1^MUT^) cells of CPRF patients and healthy fathers (two biological replicates for each condition). A comparison using principal component analysis (PCA) revealed a clear separation of the patient and healthy samples (Extended Data Fig. 1b). Differential chromatin accessibility analysis identified 16,425 peaks (from a total of 96,467 peaks) that were evenly distributed between less and more accessible in LSD1^MUT^ vs LSD1^WT^ (FDR < 0.05, Fig. 1b). More accessible regions in LSD1^MUT^ are enriched for active and bivalent/poised transcription start sites (+/- 3kb from TSS, 24% of peaks in LSD1^MUT^ vs. 6.5% of peaks in LSD1^WT^) and GC-rich promoter motifs (HOMER motif enrichment tool). In contrast, less accessible regions in LSD1^MUT^ are enriched for active and bivalent enhancers but depleted for the TSS, compared to the background peaks (Extended Data Fig. 1c). These results suggest a broadly impaired chromatin regulatory landscape in LSD1^MUT^.

**Figure 1.**
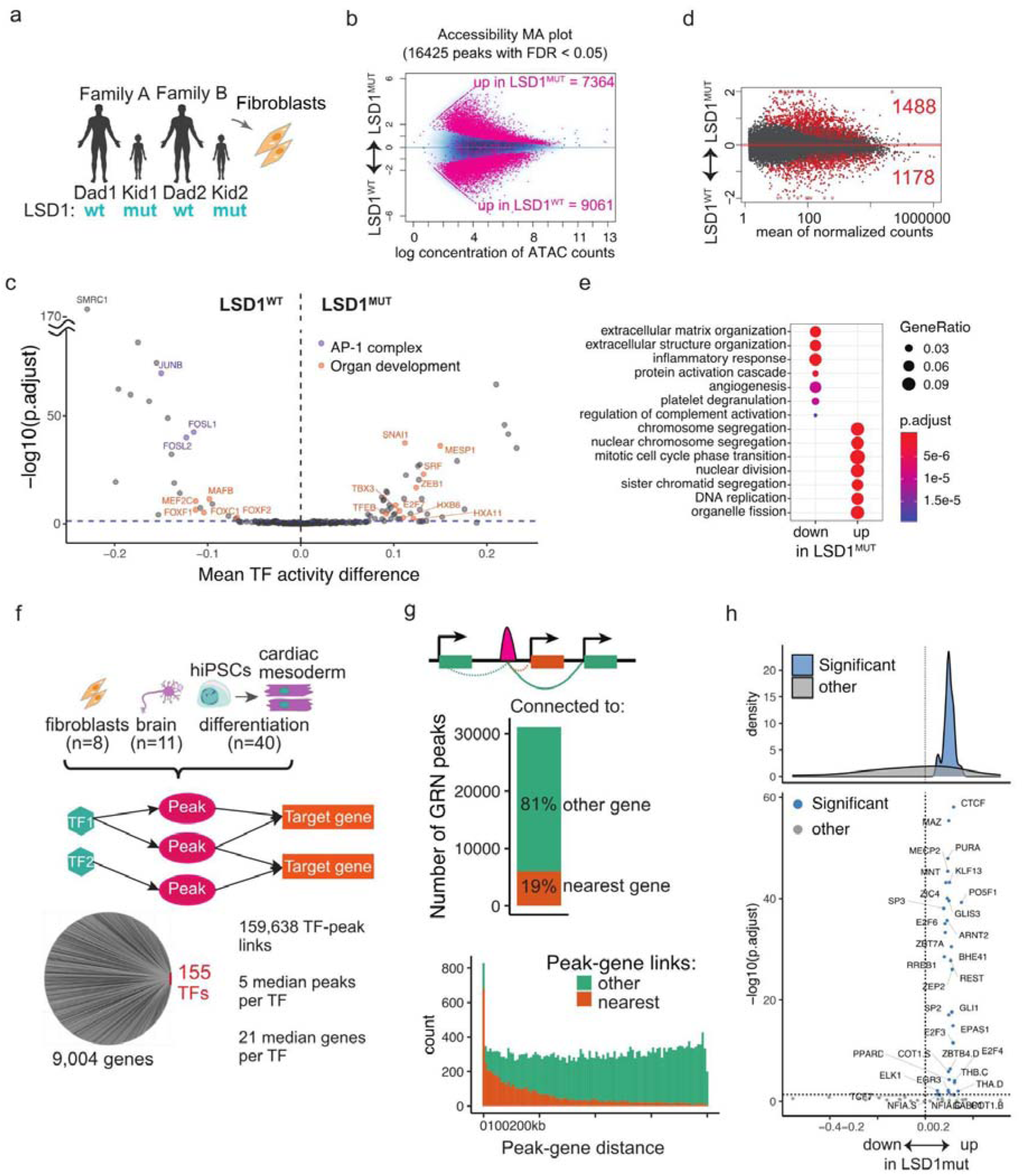
a) Schematic of the experimental set-up. b) Differential accessibility analysis with MA plot showing the concentration of accessible fragments in the ATAC-seq peaks against log2 fold change of peak accessibility in LSD1^MUT^ vs. LSD1^WT^. Significantly differential peaks (FDR < 0.05) are marked in pink. For panels c,d,f: n=4 for each condition (2 families, each 2 replicates). c) Differential TF activity analysis in primary fibroblasts comparing healthy controls (LSD1^WT^) vs patients with LSD1^MUT^. TFs colored by “AP-1 complex” (purple) and GO term “Embryonic organ development” (orange). d) Differential gene expression analysis in fibroblasts with an MA plot showing average normalized gene expression values against log2 fold change in gene expression. Significantly differential genes (FDR < 0.1) are shown in red. e) Gene ontology enrichments of significantly differential genes in fibroblasts. f) Schematic of the cell types used for eGRN construction (top, also see Table 1). A hairball plot of the full eGRN with lines (representing enhancers) connecting TFs and their target genes, with eGRN statistics. g) Top schematic: solid green line but not dotted lines shows correlated peak-gene links present in the eGRN. Middle panel: percentage of eGRN peaks linked to the nearest gene or to the other gene, from the total of 96,050 peaks. Bottom panel: the distribution of distances between peak and gene in the eGRN, colored by peaks linked to the nearest gene or other gene. h) For each TF, the mean differential expression in fibroblasts (LSD1^MUT^ vs LSD1^WT^) of its target genes (=TF regulons) is shown as a point on the scatter plot, and a density plot stratified for significant (blue) and non-significant (grey) TF regulons.

**Table 1.**

| Source | # samples | Type | Link |
|---|---|---|---|
| This study | 8 | Fibroblasts |  |
| Snyder/Murry lab | 40 | hiPSC, mesoderm, cardiac mesoderm, cardiomyocyte | https://www.ncbi.nlm.nih.gov/geo/query/acc.cgi?acc=GSE106690; https://www.ncbi.nlm.nih.gov/geo/query/acc.cgi?acc=GSE85332 |
| PsychENCODE | 11 | Brain | https://www.synapse.org/#!Synapse:syn7062404<br>https://www.synapse.org/#!Synapse:syn7349497 |

To examine the involvement of specific transcription factor (TF) binding associated with changes in chromatin accessibility between LSD1^MUT^ and LSD1^WT^, we applied diffTF analysis^14^, which quantifies genome-wide differential TF motif accessibility as “TF activity,” with increased TF motif accessibility corresponding to higher TF activity, and decreased accessibility to lower activity. Comparing the TF activities of LSD1^MUT^ and LSD1^WT^, and excluding age-related TFs to minimize the effect of the donors’ age (see Methods), identified 68 significantly changed TFs (Fig. 1c). The 44 TFs with increased activity in LSD1^MUT^ include developmental factors (Tbx3, Mesp1, Hoxb6/a11) and the epithelial-to-mesenchymal transition (EMT) TFs Zeb1 and Snai1, whereas several TFs with decreased activity include the AP-1 complex, which controls responses to various stimuli. Protein-protein interaction data (STRING database) showed greater interconnection between differentially active TFs than expected by chance (average node degree is 3.85 compared to 1.5 in a network of non-significant TFs and protein-protein interaction enrichment p-value is < 1e-16), suggesting that genome-wide chromatin accessibility differences in LSD1^MUT^ are associated with a group of functionally related TFs.

### LSD1^MUT^ induced changes in promoter accessibility and expression

Having identified TFs (inferred by motif) that are mis-regulated in LSD1^MUT^, we sought to determine transcriptional changes in fibroblasts. RNA-seq analysis of LSD1 mutants and controls revealed more than 2500 differentially expressed genes (Fig. 1d), of which 1488 were upregulated and 1178 downregulated in LSD1^MUT^ (DESeq2, FDR < 0.1, Fig. 1d). Up-regulated genes were enriched for cell cycle transition and DNA replication GO terms (Fig. 1e), however upregulation of cell cycle genes appeared in only one (Glu403Lys) of the two LSD1^MUT^ samples, undermining its wide relevance in CPRF syndrome. The LSD1 Glu403Lys mutant, but not Tyr785His mutant, showed a strong impairment in histone H3 binding^13^, which could decrease localization of repressive co-factors, resulting in upregulation of target genes. Down-regulated genes in LSD1^MUT^ were enriched for the extracellular matrix, inflammatory response, and angiogenesis GO terms, consistent with the role of extracellular matrix in the EMT and impairment of the repressive EMT TFs Zeb1 and Snai1 identified in the diffTF analysis.

To assess the relationships between chromatin accessibility and gene expression changes in LSD1^MUT^ fibroblasts, we linked each ATAC-seq peak at promoter region to the gene and analyzed these peak-gene links (Ext. Data Fig. 1e). We observed a positive correlation between significant differential chromatin accessibility and gene expression (Pearson’s coefficient = 0.42). Top correlated links appear in RBM46, DDX43, and TLR4 genes. Yet, the numbers of linked cases (n=486) were low and limited to the promoter peak-gene links, indicating that a more robust approach is required to analyze the functional interactions between changes in chromatin accessibility and gene expression.

### TF-gene modules affected by LSD1^MUT^

To dissect the link between changes in chromatin accessibility at regulatory elements and gene expression beyond simply proximity, and to connect changes in TF activity with downstream effects on target gene expression, we constructed an enhancer-mediated gene regulatory network (eGRN) based on co-variation across samples with paired RNA-seq and chromatin activity readout ^15, 16^ such as ATAC-seq data using GRaNIE software ^17^. We calculate correlations across samples between TF expression and ATAC-seq signal at their binding motifs (TF-to-peak correlations), and between ATAC-seq peaks and RNA expression of nearby genes (peak-to-gene correlations, see Methods) to connect the genes to their putative regulatory elements and TFs. To obtain the TF-enhancer-gene links in the cell types relevant for the LSD1^MUT^ and CPRF syndrome phenotype, we used published datasets from healthy brain samples, two independent studies on a differentiation time course spanning pluripotent stem cells, via mesoderm, to cardiomyocytes, in addition to our fibroblast data, containing paired ATAC- and RNA-seq data (Table 1 and Fig. 1f). After filtering for significant TF-to-peak and peak-to-gene connections, we obtained a network of 155 TFs linked to 9,004 unique genes via 31,146 regulatory elements (Fig. 1f and Methods). This large network represents the global set of all TF-target gene connections in healthy cell types. In this eGRN, only 19% of peaks were linked to the nearest gene, consistent with our previous study reporting <20% of linked peaks^15^. The majority of peaks were linked to distal genes rather than the nearest gene within 500 kb of the TSS (Fig. 1g), implying greater regulatory network complexity and the relevance of using cell type specific ATAC- and RNA-seq data to derive peak-gene connections.

We defined the genes connected to each TF in the eGRN as TF regulons (TF-gene modules). For each TF, we assessed whether its regulon activity, i.e. the distribution of aggregated expression changes of all TF target genes compared in LSD1^MUT^ vs. LSD1^WT^ fibroblasts, was significantly different from the expression changes in all other genes (see Methods and Extended Data Fig. 2a). Globally the distribution of gene expression of all 9,004 genes retained in the eGRN was not affected by LSD1^MUT^ (mean log2 fold changes are centered around 0, Extended Data Fig. 2b), however we found 34 TFs whose regulons’ expression (log2 fold change in LSD1^MUT^ vs. LSD1^WT^) in fibroblasts deviated significantly from the background (Fig. 1h). All regulons were upregulated in LSD1^MUT^ fibroblasts, consistent with an overall de-repression of genes due to reduced LSD1 function and perturbed repressor TF activity, as seen in the de-repression of the REST, MECP2 and ZBTB4 regulons (Fig. 1h). The differential activity of this set of TFs, based on ATAC-seq data, correlates with the activity of their regulons from the eGRN (pearson correlation: 0.4, Extended Data Fig.2c), confirming the direct effects of LSD1^MUT^-mediated TF motif accessibility disruption on TF target gene expression. Our eGRN therefore identified a specific set of TFs (inferred by motif) that likely mediate the changes in gene expression in LSD1^MUT^ fibroblasts, with many of these changes being upregulation of genes.

### Gene expression during lineage differentiation in LSD1^MUT^

The role of LSD1 in disabling enhancer regions through the removal of H3K4me1/2 marks is critical during cellular differentiation, when the enhancer landscape has to be re-established^18^. To study the differentiation of patient-derived cells, we reprogrammed patient- and control-derived primary fibroblasts from the two families into human induced pluripotent stem cells (hiPSCs). All hiPSC clonal lines (n= 3 from family A and n = 2 from B, LSD1^WT^; n = 2 from family A and n = 2 from B, LSD1^MUT^) show normal chromosome numbers, intense staining of pluripotency markers (e.g., Oct4, SSEA-4, TRA-1-60), and high expression levels of pluripotency marker genes (Extended Data Fig.3a). We differentiated all nine hiPSC lines into three main embryonic lineages: endoderm, mesoderm, and the neuronal lineage, and analyzed the effects of LSD1^MUT^ on gene expression (Fig. 2a-b). In the pluripotent state, we observed limited changes in gene expression between LSD1^MUT^ and LSD1^WT^ (58 differentially expressed genes, Fig. 2c) partially due to the high variability between the hiPSC lines (Extended Data Fig. 3b), reminiscent of other reprogrammed hiPSC studies^19^. However, during differentiation into the three lineages, LSD1^MUT^ had a substantial impact on gene expression. Over 4700 genes were differentially expressed in endoderm, over 1600 in the neuronal lineage, and over 700 in mesoderm (FDR < 0.05, Fig. 2c), consistent with the phenotype of the affected patients who show symptoms related to all lineages (e.g., developmental delay, muscle hypotonia, distinctive facial features). While most of the differential genes were mis-regulated in only one of the cell lineages, a larger set (596 genes) were mis-regulated in both endoderm and neuronal lineages, and 87 genes were mis-regulated in all three lineages.

**Figure 2.**
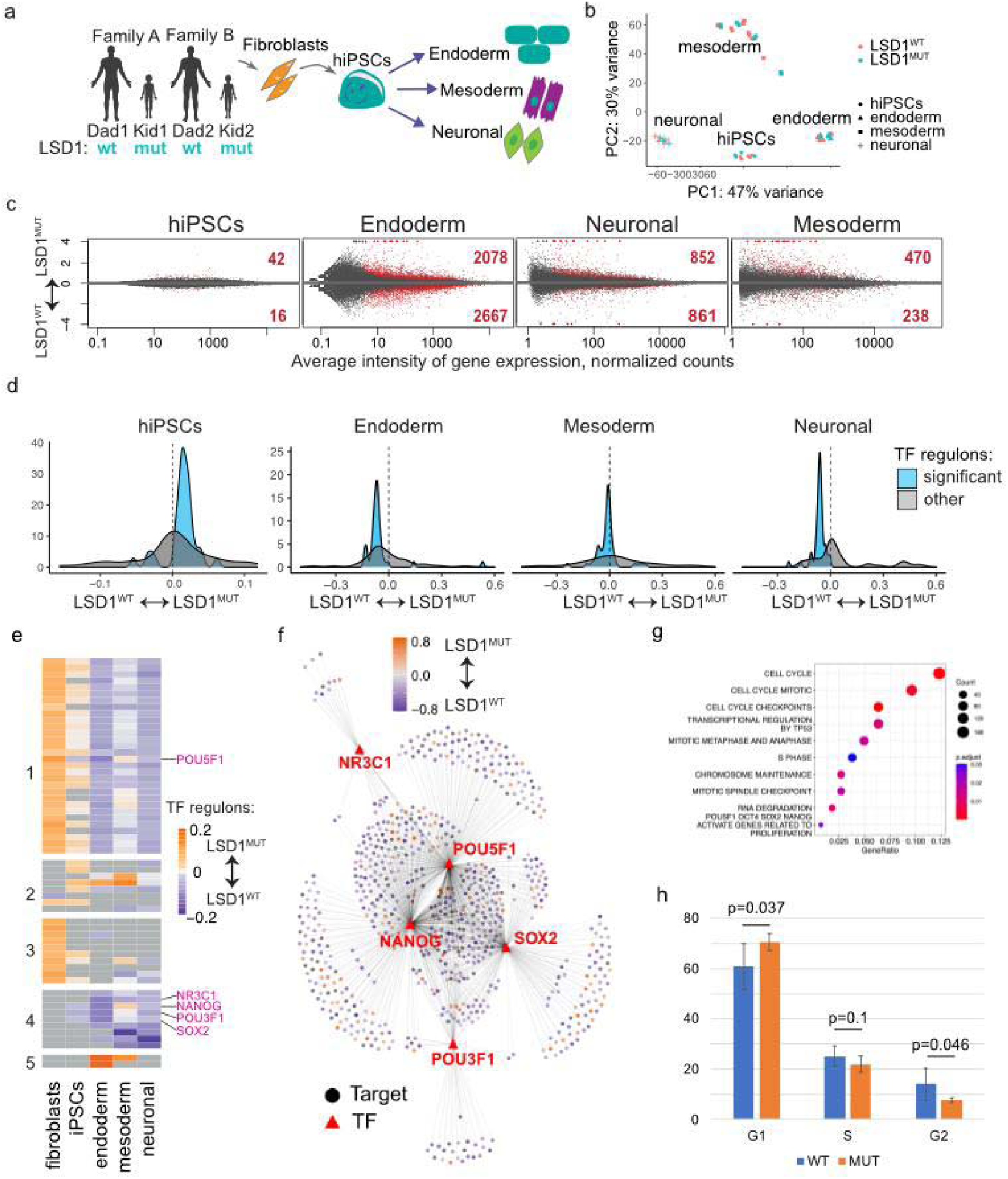
eGRN analysis identifies aberrant gene repression by LSD1^MUT^ during the early differentiation of cell lineages. a) Reprogramming of primary fibroblasts into hiPSCs and subsequent differentiation into three main embryonic lineages. b) Principal component analysis (PCA) plot of RNA-seq data in hiPSCs and derived from them endoderm, neuronal and mesoderm lineages. Principal components 1 and 2 are shown together with the variance they explain. c) Differential expression analysis of LSD1^MUT^ effects on hiPSCs and early differentiation in three embryonic lineages. n=8 for LSD1^MUT^ and n=10 for LSD1^WT^. d) Density plots of TF regulons expression in LSD1^MUT^ hiPSCs and derived from them endoderm, mesoderm and neuronal lineages. Significant TF targets are upregulated in hiPSCs but downregulated in lineages. e) Heatmap of a GRN subnetwork showing average expression changes of TF regulons in LSD1^MUT^ across cell types. This subnetwork includes 59 TFs whose targets’ expression is significantly changed in LSD1^MUT^ cell lineages. f) Graph representation of an eGRN subset of the 5 TFs, whose regulons were most affected in endoderm differentiation. Only significantly differential TF targets (circles) are shown, colored by their expression log2 fold change in LSD1^MUT^ vs. LSD1^WT^. g) Gene ontology enrichments of TF regulons from the eGRN subset in panel f. h) Cell cycle analysis of the endoderm cells (LSD1^MUT^ and LSD1^WT^)

We observed that genes downregulated in endoderm and neuronal lineages were enriched in cell cycle related GO terms (DNA replication, sister chromatid segregation, Extended Data Fig. 3c). Unlike the increase in cell cycle genes observed in one of the two LSD1^MUT^ individuals in primary fibroblasts, the downregulation of cell cycle related genes across differentiating lineages occurred in both LSD1^MUT^ samples, implying more relevance to CPRF syndrome. Differentiation efficiency and the overall cell morphology in each lineage were not affected by LSD1^MUT^, consistent with comparable expression levels of lineage-specific marker genes (Extended Data Fig. 3d) and suggesting that the decreased expression of cell cycle genes is not caused by developmental defects.

The downregulation of genes in differentiated endoderm and neuronal cells observed in LSD1^MUT^ could result from reduced activation of up-regulated genes during differentiation or increased repression of down-regulated genes. To distinguish between these two possibilities, we compared gene expression changes in LSD1^MUT^ and LSD1^WT^ endoderm split into up- and down-regulated genes relative to expression levels in hiPSCs (Extended Data Fig. 3e). This analysis revealed that genes whose expression decreased in LSD1^WT^ endoderm compared to hiPSCs were more strongly downregulated in LSD1^MUT^ endoderm. Given the reduced cell proliferation capacity when transitioning from hiPSCs to defined cell lineages, genes in these categories decreased in expression, but the decrease was stronger in LSD1^MUT^.

### eGRN TF regulons downregulated in LSD1^MUT^ at early lineage differentiation

Our eGRN identified a specific set of TFs (inferred by motif) whose regulons were mis-regulated in LSD1^MUT^ fibroblasts. To determine the effects of LSD1^MUT^ on TF-gene pairs in the other cell types, we performed the TF regulon analysis in LSD1^MUT^ vs LSD1^WT^ hiPSCs and hiPSC-derived cell lineages. Similar to the fibroblast data, most TF regulons were upregulated in LSD1^MUT^ hiPSCs (Fig. 2e), consistent with the LSD1 repressive function. However, during early cell differentiation (from hiPSCs to endoderm, mesoderm, and neuronal lineages) the majority of TF regulons were downregulated in LSD1^MUT^ (Fig. 2e). To analyze the regulons across different lineages, we focused on TFs that were not differentially expressed following differentiation (FDR > 0.1 and absolute log2FC < 0.58) but whose regulon activity was significantly altered in LSD1^MUT^. This identified 59 TFs, which were then grouped into five classes based on unsupervised clustering of their regulon activity across cell types (Fig. 2e). Cluster 1 contains TFs with upregulated regulon activity in fibroblasts and hiPSCs, and downregulated in endoderm, mesoderm, and neuronal lineage in LSD1^MUT^ vs LSD1^WT^. Clusters 2 and 5 TF regulons are unchanged in fibroblasts, with individual TFs upregulated in either hiPSCs or one of the differentiation lineages. Cluster 3 TF regulons are almost exclusively upregulated in fibroblasts, with little changes in other cell types, whereas cluster 4 TF regulons are not altered in fibroblasts but are downregulated in the differentiated cell lineages. Thus, all TF regulons are repressed by LSD1^MUT^ in the three differentiating lineages, whereas TF regulons are predominantly upregulated by LSD1^MUT^ in fibroblasts and hiPSCs (Fig. 1h and Fig. 2d-e). Upregulated TF regulons are separated into three distinct sets (clusters 1-3), suggesting that LSD1^MUT^ affects TF activity in a cell type-specific manner. The effect size of the repression is the strongest in cluster 4 and overall less substantial in the mesoderm (Fig. 2e). Functional enrichment analysis among TF regulons in five Clusters revealed significant overrepresentation of cell proliferation related categories such as translation and cell cycle in clusters 1 and 4 (Extended Data Fig. 3f).

To examine specific TFs (inferred by motif) and their regulons in detail, we selected the top 5 TFs whose regulons were significantly altered in endoderm by LSD1^MUT^ (Fig. 2f). Three of the five top TFs are the core pluripotency factors: Nanog, Oct4 and Sox2. Previous work has shown that LSD1 is required for decommissioning of active enhancers during mouse ESC differentiation ^18^. In contrast to the previous work, LSD1^MUT^ led to over-repression of the pluripotency-related regulons in differentiating endoderm and neuronal lineages, and to some extent in the mesoderm (Fig. 2e-f). The 5 top TFs are linked to 710 genes differentially expressed in LSD1^MUT^ endoderm, with the majority of these genes (414) are regulated by at least 2 TFs, in accordance with the overlapping function of the core pluripotency factors. Gene targets of the top 5 TFs are overrepresented in pathways related to highly proliferative cells (cell cycle, mitosis, Oct4/Nanog/Sox2 targets) (Fig. 2g). We confirmed the changes in cell cycle in LSD1^MUT^ endoderm cells by FACS analysis, which identified reduction of LSD1^MUT^ cells in S-phase (although not significant) and G2 phase, and significant increase in the proportion of resting G1 cells (Fig. 2h). These results indicate that during early lineage differentiation pluripotency TF regulons get repressed and LSD1^MUT^ further enhance this repression, contrary to the expected de-repression from a loss of function phenotype of LSD1.

Additionally, distinct TF regulon clusters could be associated with specific genetic risk factors. To investigate if regulatory regions associated with TF regulon clusters 1-5 are enriched for common genetic traits, we performed linkage disequilibrium score regression (LDSC) analysis^20^ using 442 GWAS catalogue traits (ExtDataFig. 3g and Methods). The TF regulons of cluster 1 are enriched for immune, adipose, blood and brain-related GWAS traits (osteoarthritis, body mass index, cholesterol measurement, leukocytes, epilepsy), suggesting that the inverse TF regulon expression patterns observed in fibroblasts/hiPSCs and early differentiation are associated with multiple cell types. The TF regulons of clusters 2 and 3 are enriched in one psychiatric/brain related trait each (loneliness and feeling miserable measurements), preventing drawing sound conclusions from the single traits. The TF regulons of cluster 4, which are down-regulated in early differentiation, are enriched in GWAS traits related to multiple organs (allergy, body fat, snoring, FEV/FVC ratio for lung function), consistent with the multi-organ phenotypes of the CPRF patients and indicating the relevance of using hiPSC-derived cell differentiation approaches for *in vitro* disease modeling.

### LSD1^MUT^ increases HDAC activity and repression of target genes

To understand how TF regulons are being repressed in LSD1^MUT^ during early differentiation, we considered two possibilities: a dosage effect and a mutation-specific effect (Fig. 3a). A dosage effect was observed in heterozygous LSD1^MUT^ fibroblasts and hiPSCs, where TF regulons were upregulated due to the reduction in LSD1-mediated repression, consistent with the previous work ^18^. The stronger repression of TF regulons seen during early differentiation in LSD1^MUT^ could be caused indirectly, e.g. by overexpression of repressive TFs. Indeed, 14 repressor TFs are upregulated in endoderm, mesoderm, or the neuronal lineage but none of them significantly increased in all three cell types (Extended Data Fig.4a), whereas the global downregulation of TF regulons is found in all three lineages, suggesting that overexpression of repressors does not fully explain the molecular phenotype. A mutation-specific effect is suggested by a previous biochemical study of CPRF patients’ mutations^13^, which has shown that the LSD1^MUT^ is still able to interact with its repressive co-factors (HDACs and the Co-REST complex) and, to some extent, histone substrates. Because many LSD1 co-factors have a repressive role, their increased recruitment and/or activity in LSD1^MUT^ cells could lead to the decrease in histone acetylation and observed over-repression of TF regulons (Fig. 3a). Indeed, a previous in vitro study has shown that LSD1 and HDAC1 are enzymatically and structurally coupled within the Co-REST complex ^21^, suggesting that catalytically inactive LSD1^MUT^ protein could affect repressive activity of HDAC1.

**Figure 3.**
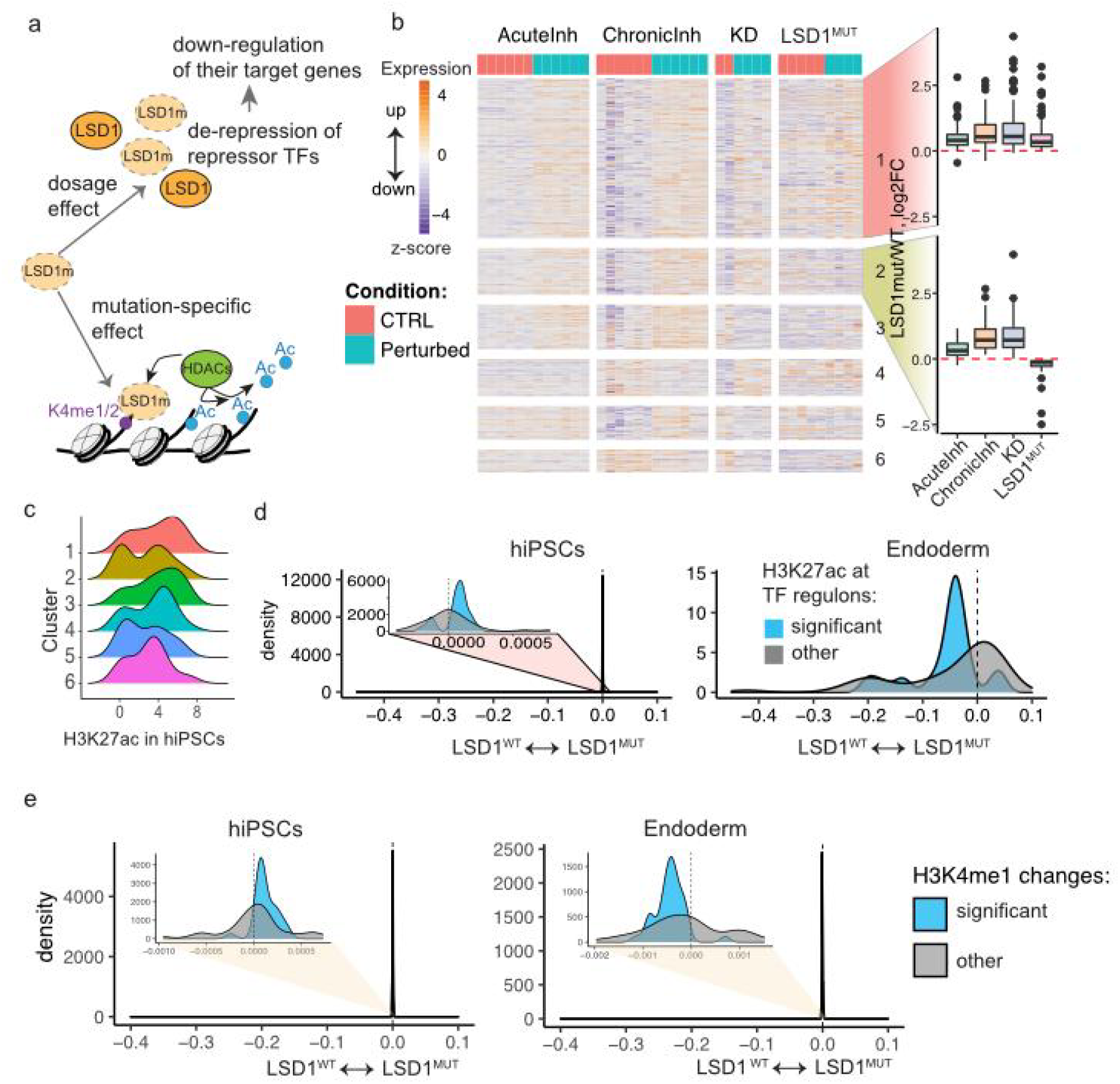
Increased HDAC activity leads to over-repression of TF targets in LSD1^MUT^. a) Scheme of dosage and mutation-specific effects in LSD1^MUT^. b) Heatmap of gene expression changes in the control and perturbed hiPSCs (acute or chronic chemical inhibition of LSD1, knockdown of LSD1 by shRNA and CPRF patient-derived cells). Normalized expression values are shown as z-scores within each treatment batch. Boxplots of log2 fold changes between perturbed and control cells in each condition are shown on the right for the two first clusters. c) Density plots of H3K27ac signal in the LSD1^WT^ iPSCs at the regulatory regions linked to the genes in the heatmap on panel B. Cluster 2 shows strong bimodal distribution of low- and high-K27ac regions. d) H3K27ac ChIP-seq signal at the TF regulon-linked enhancers, X-axis shows changes between LSD1mut and LSD1wt lines. Inset plot showing marginal effect of H3K27ac in hiPSCs. e) H3K4me1 ChIP-seq signal at the TF regulon enhancers showed very small differences between the LSD1^MUT^ and LSD1^WT^ samples (0.5-1%, see inset plot).

To further characterize the dosage and mutation-specific effects we used RNA-seq to analyze normal hiPSCs treated with either a chemical inhibitor of LSD1 (acutely, for 24h, or chronically, for 7 days) or shRNA to knock-down LSD1 (see Methods). Comparing gene expression changes arising from these perturbations (reduced dosage of LSD1) with the LSD1^MUT^ data from reprogrammed hiPSCs (mutation-specific condition) allowed the identification of dosage and mutation-specific effects (Fig. 3b). Genes that were differentially expressed in any of these conditions grouped into 6 main clusters, with most genes (cluster 1, n=133) being upregulated, given the loss of LSD1 repressive functions (Fig. 3b). The transcriptional repressor Zeb1 was among the upregulated genes, its increased expression potentially resulting in stronger downregulation of its targets and contributing to the phenotype we observed during hiPSCs differentiation (Fig. 2d). The second cluster (n=39 genes) contained genes that were downregulated only in LSD1^MUT^, suggesting mutation-specific regulation. Assessing HDAC activity based on its substrate H3K27ac abundance in hiPSCs (see Methods), we found genes in the second cluster were linked to chromatin regions with less H3K27ac signal (Fig. 3c). These results suggest that there is increased HDAC activity in the LSD1^MUT^ cells and not in the cells with reduced dosage of LSD1.

Next we tested whether the abundance of HDAC substrate H3K27ac, or LSD1 substrate H3K4me1, are also reduced at the enhancers associated with downregulated regulons in LSD1^MUT^ endoderm cells and performed ChIP-seq experiment in patient-derived samples (Fig. 3d-e). Indeed, H3K27ac levels were decreased specifically at the significantly downregulated regulons in LSD1^MUT^ endoderm cells, and not in all regulons, and neither in hiPSC cells (Fig. 3d, negligible size effect in hiPSCs). H3K4me1 signal was nearly unaffected in LSD1^MUT^ hiPSCs and endoderm (Fig. 3e), pointing towards the stronger effect of LSD1^MUT^ on HDAC activity and not on H3K4me-dependent chromatin impairment.

### HDAC repression in LSD1^MUT^ reverses altered gene expression

Given the connection between decreased gene expression in LSD1^MUT^ and increased HDAC activity, we hypothesized that chemical inhibition of HDAC activity might rescue repression of the TF regulons during differentiation. To test this, we differentiated hiPSC lines (n= 5, LSD1^WT^; n = 4, LSD1^MUT^) into endoderm, treating differentiating cells with the HDAC inhibitor valproic acid (VPA), 400µM, or vehicle control, and measured changes in gene expression (LSD1^MUT^ vs. LSD1^WT^ cells). VPA, approved by the FDA for the treatment of epilepsy and mood disorders^22^, inhibits both class I and class IIa HDACs ^23–25^, increasing the levels of acetylated histones (Extended Data Fig.4b). In the untreated LSD1^MUT^ endoderm we identified 1170 differentially expressed genes. VPA treatment rescued expression of nearly all of these genes, with only 21 remaining significantly differentially expressed (Fig. 4d). Most of the genes that were downregulated in LSD1^MUT^ and rescued by VPA treatment were expressed at levels close to LSD1^WT^ levels (Fig. 4e).

**Figure 4.**
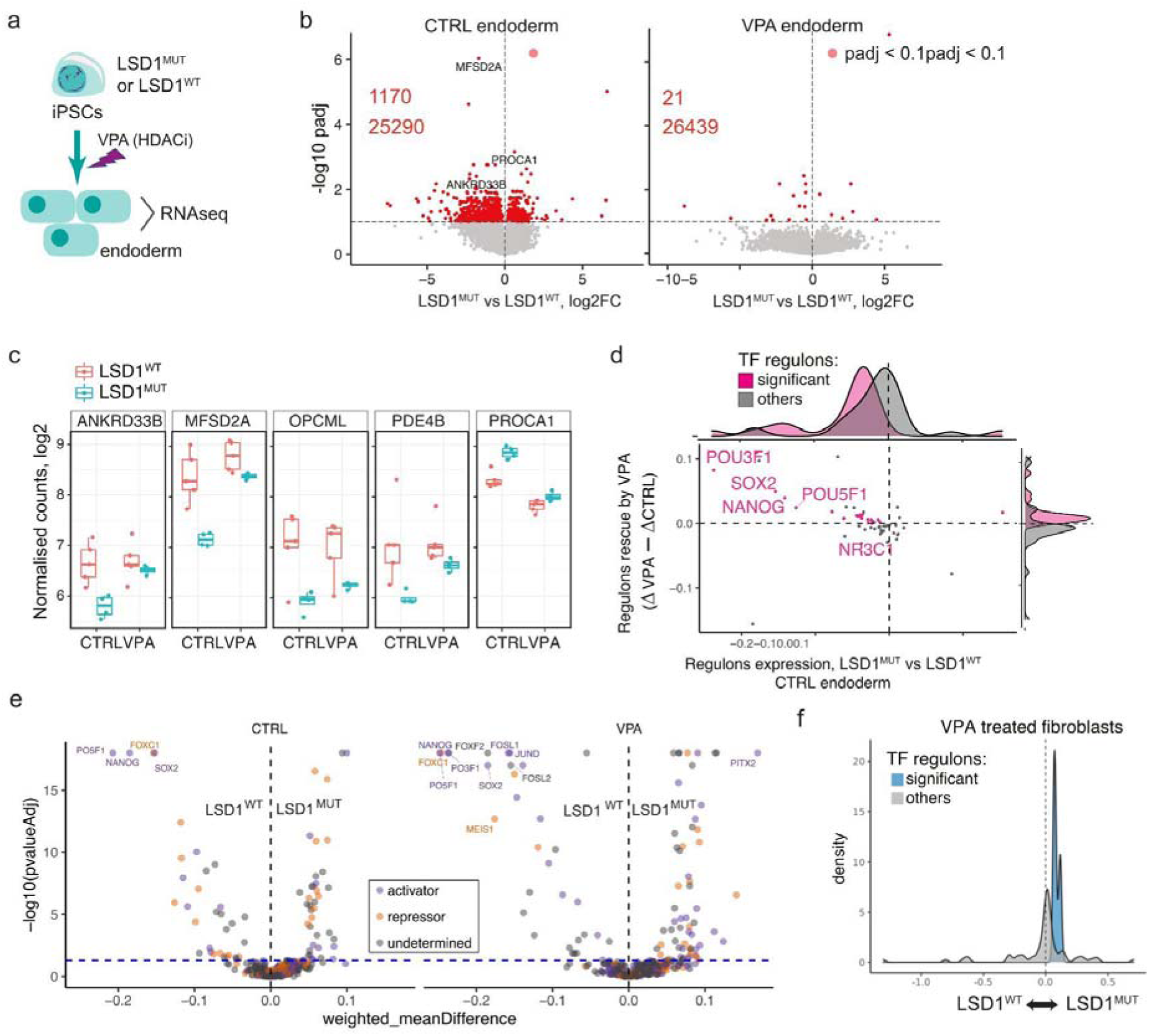
Valproic acid (VPA) treatment rescues the over-repressive TF regulons in the LSD1^MUT^ endoderm. a) Scheme of the VPA treatment experiment: CPRF patient-derived hiPSCs were differentiated within 48h to endoderm in the presence of HDAC-inhibitor VPA and gene expression was measured by RNA-seq. b) Volcano plots show the distribution of gene expression changes in LSD1^MUT^ vs LSD1^WT^ control and VPA-treated endoderm. Significantly differential genes (padj < 0.1) are highlighted in red. n=4 for LSD1^MUT^ and n=5 for LSD1^WT^ (same for panel c). c) Expression counts of selected top differentially expressed genes in control and VPA treated endoderm. d) Scatter plot of the significantly affected by LSD1^MUT^ TF regulons in control endoderm, showing the difference □ in TF regulon expression between LSD1^MUT^ and LSD1^WT^ control endoderm (X-axis) against the size effect of the VPA-mediated rescue of TF regulon expression (□ in VPA - □ in control endoderm). e) *Differential TF activity analysis in patient-derived endoderm (CTRL or VPA-treated) comparing healthy controls (LSD1^WT^) vs patients with LSD1^MUT^. Top differentially active TFs are labeled, colors correspond to predicted TF role as activators or repressors. f) Density plots of TF regulons expression in LSD1^MUT^ fibroblasts treated with VPA. Significantly affected regulons (in blue) remain upregulated as in Fig.1h despite VPA treatment*.

To investigate the basis for the highly effective VPA-mediated rescue, we examined the TF regulons and identified the top 30 TF motifs (Extended Data Fig. 4c) that were linked to the genes rescued by VPA. Unsupervised k-means clustering of these TFs identified two enriched groups: regulation of pluripotent stem cells (MECP2, SOX2, OCT4, NANOG) and mitotic cell cycle regulation (SP1/3, E2F1/6, TFDP1). This result suggests that VPA treatment is rescuing the TF regulons related to stemness and cell proliferation, impaired by LSD1^MUT^.

To assess the global effects of VPA treatment on TF regulons, we compared the TF regulon expression changes in untreated and VPA-treated LSD1^MUT^ endoderm. We observed upregulation of TF regulons and partial rescue of the LSD1^MUT^ effects on global gene expression (although not significant, p=0.18, Extended Data Fig. 4d). TF regulons whose expression was most rescued by VPA significantly overlapped with the most affected TFs in endoderm differentiation (Fig. 3e), including the core pluripotency TFs (Nanog, Sox2 and Pou5f1/Oct4 Fig. 2e-f and 4d).

We further sought to identify whether the rescue effect of the VPA treatment is mediated via chromatin and performed ATAC-seq experiment on VPA-treated and control endoderm cells. TF activity analysis using diffTF showed reduced activity of the core pluripotency TFs Oct4, Sox2 and Nanog as the most affected TFs in control endoderm (Fig. 4e), in accordance with the TF regulon expression analysis of the GRN. Notably, VPA treatment did not rescue the activity of these TFs (Fig. 4e), indicating that HDAC inhibition did not impact chromatin accessibility of the TF binding sites and the rescue effect of the VPA treatment is not mediated by chromatin decompaction of the TF binding sites.

While the rescue effect of TF regulons expression impaired by LSD1^MUT^ is a promising finding with potential applications for the affected patients, so far we have shown it only during early cell differentiation, making it less applicable to young/adult individuals. We therefore tested whether VPA treatment affects differentiated cells and performed RNA-seq analysis on VPA-treated patient-derived fibroblasts. TF regulon expression of significantly affected TFs was upregulated in fibroblasts (Fig. 1h) and VPA treatment did not impact this upregulation (Fig. 4f), arguing against any strong effect of the VPA on terminally differentiated cells.

## Discussion

De novo point mutations in the histone lysine demethylase LSD1 cause a rare neurodevelopmental disorder, CPRF. However, the molecular mechanisms underlying the disorder are not known. Using CPRF patient-derived primary fibroblasts, hiPSCs and differentiated cell lineages we studied the effects of two different LSD1 mutations on epigenomic and transcriptional landscape, as well as on TF-target gene interactions. We found that in defined cell types (primary fibroblasts and hiPSCs) LSD1^MUT^ leads to the overall upregulation of TF regulons, consistent with a reduction in LSD1 repressive function. In contrast, during the early differentiation of hiPSCs into endoderm, mesoderm and neuronal lineages LSD1^MUT^ had a dominant effect, promoting increased repression of TF regulons, mostly affecting cell proliferation related genes. We have identified HDACs as mediators of the LSD1^MUT^ repressive phenotype and have shown that inhibition of HDAC activity using a chemical compound, VPA, rescues the repressive phenotype in LSD1^MUT^ endoderm differentiation.

LSD1^MUT^ caused changes in activity of a specific set of TFs, as well as expression of their regulons in all analyzed cell types. However, the set of TFs and the direction of the expression changes of their regulons varied between cell types. Such cell type specific differences could arise from the activity of distinct enhancers in particular cell states^26^, or from relocation of LSD1 to different enhancers during cell state transitions. Moreover, interactions of LSD1 with its co-factors and TFs can vary between cell types, for example, through tissue-specific TFs^27^, introducing another source of variability in the molecular phenotype of LSD1^MUT^. Sensitivity of a particular set of TFs to LSD1^MUT^ might reflect the susceptibility of TF binding sites to the chromatin environment altered by LSD1^MUT^.

Our eGRN identified enhancer-mediated TF-target gene connections directly affected by LSD1^MUT^, facilitating the dissection of LSD1 effects into dosage loss of function of LSD1 and dominant effects of LSD1^MUT^. Thus, CPRF molecular pathology cannot be fully dissected using the only loss of function model systems due to substantial mutation-specific transcriptional effects. LSD1^MUT^ has a dominant repressive effect on TF regulon expression during cell differentiation. We show that this effect is linked to HDACs, known LSD1 co-factors. HDAC inhibitors were previously found to rescue a deficiency in KMT2D, an H3K4 methyltransferase, by balancing the lack of active H3K4 marks^28^, whereas in this work, the rescue of a repressor LSD1 impairment is achieved by inhibiting another repressor (HDAC), suggesting that HDACs have higher activity or recruitment of in the presence of LSD1^MUT^ protein. In the fibroblast and hiPSC cell states we did not observe increased repression of target genes, suggesting this molecular phenotype might be transient and compensated for later by the wild-type copy of LSD1. Nevertheless, the defects in early cell differentiation could have profound consequences on tissue formation and organism development.

The selectivity of HDAC effects on certain enhancers can be explained by different “basal” levels of the histone marks at these enhancers. For example, it has been reported that LSD1 and HDAC within the CoREST complex cannot bind their nucleosome substrates and their enzymatic products simultaneously^21^, thus the levels of H3K4me1/2 and H3K27ac could influence the activity of the HDAC and LSD1, respectively. Our ChIP-seq experiments in endoderm did not show significant differences in basal levels of H3K27ac or H3K4me1 between “affected” enhancers and other enhancers (data not shown). Instead, in iPSCs, affected enhancers exhibited significantly higher basal H3K4me1 and H3K27ac, marks of active enhancers known to be bound by LSD1 already in stem cells^18^, which suggests that these regions would be more affected by LSD1 mutations upon differentiation.

Since acetylated enhancers are often associated with p300 acetyltransferase, which counteracts HDAC activity^29^, this may explain the lack of HDAC-mediated repression in the iPSCs and instead a delayed effect on the expression of the target genes later during differentiation to endoderm.

We hypothesize that the dominant LSD1^MUT^ effect prevails over haploinsufficiency specifically during cell state transitions, such as early stages of stem cell differentiation. LSD1 demethylation activity is inhibited in the presence of acetylated histones^30^, and during stem cell differentiation H3K27ac levels decrease at enhancers of proliferation-related genes^18^, enabling LSD1 to demethylate these regions. The inability of the LSD1^MUT^ protein to demethylate H3K4me1/2 leads to aberrant retention of this modification on chromatin, and instead of diffusion of LSD1 after productive demethylation, further attracts LSD1^MUT^ protein. This could potentially lead to longer residence times for the LSD1^MUT^-HDAC complex at its target regions, causing even stronger repression of TF regulons by HDAC. This effect is likely transient and compensated for by the functional LSD1^WT^ copy after cells reached their differentiated state (e.g. in fibroblasts). In the differentiated cells LSD1^MUT^ resembles haploinsufficiency, with the upregulation of TF regulons likely caused by an overall increase in poised enhancers marked by H3K4me1, due to the lack of demethylation activity in LSD1^MUT^.

Overall, we show that LSD1^MUT^ is linked to misregulation of the global chromatin regulatory landscape, which affects motif accessibility of a specific set of TFs and the expression of genes related to cell proliferation, differentiation and development. We also identified a group of TFs whose activity was impaired in a cell-type specific manner by LSD1^MUT^ and have shown that an FDA-approved drug, VPA, can rescue TF regulon expression defects caused by LSD1^MUT^ during cell differentiation, suggesting VPA might provide a potential therapeutic avenue to explore for treatment of CPRF syndrome patients. Given that LSD1 levels are often increased in various cancers^11,30–32^, our discovery of the misregulation of LSD1-HDAC activity in regulatory regions could also contribute to the improvement of anti-cancer therapies.

## Methods

### Data availability

RNA-seq and ATAC-seq raw data for patient-derived human samples have been submitted and are currently processed by the European GenomePhenome Archive. The accession numbers will be communicated to the editor as soon as we receive them. Restrictions apply to the availability of the raw data (deposition in a controlled access repository) due to reasons of sensitivity of the human samples.

Processed ATAC-seq samples (bigwig genome coverage files, hg19 version) are deposited at EMBL OwnCloud and can be accessed via following link: https://oc.embl.de/index.php/s/fCAO7QCm2MtOmJx.

### Code availability

Custom code used to perform the analyses is available from github: https://github.com/dariabunina/LSD1_mutation_study.

### Cell lines and culture conditions

Primary skin fibroblasts were obtained from two male pediatric patients and their healthy fathers under a protocol approved by the Institutional Review Board at Translational Genomics Research Institute (WIRB; study number 20120789). We have obtained informed consent from all participants. Fibroblasts were cultured at 37 C under 5% CO2 in DMEM media supplemented with 10% fetal bovine serum. hiPSCs were generated from primary fibroblasts using Sendai virus reprogramming kit (ThermoFisher Scientific), yielding 2-3 different clonal lines for each individual. hiPSCs were maintained in Essential 8 (E8) media (LifeTech) on vitronectin coated plates following the manufacturer’s instructions. hiPSCs were passaged at least twice after thawing prior to the start of differentiation into endoderm, mesoderm and neuronal lineages. For endoderm differentiation we used a PSC Definitive Endoderm (DE) Induction Kit (Gibco, ThermoFisher Scientific Cat. No. A3062601), cells were collected after 2 days. For mesoderm differentiation we used a PSC cardiomyocyte differentiation kit (ThermoFisher Scientific Cat. No. A29212), cells were collected after 4 days. For neuronal differentiation we used a Gibco PSC neural induction medium (ThermoFisher Scientific Cat. No. A1647801) and cells were collected after 7 days. For the VPA treatment hiPSCs were differentiated to endoderm using a 2-day DE Induction Kit following the manufacturer’s instructions and from day 0 of differentiation 400uM VPA (valproic acid sodium salt 98%, Sigma Cat. # P4543-10G) in DMSO was added to cell cultures.

Healthy human hiPSC cell lines for Figure 4 were cultured under feeder-free conditions on plates coated with human-qualified Matrigel (BD Biosciences) diluted 1:40 in mTeSR-1 medium (StemCell Technologies). hiPSCs were passaged upon treatment with Accutase (Sigma) and resuspending them in mTeSR-1 medium supplemented with 5 μM Y-27632 (Sigma). Six hiPSC lines from 3 different individuals were treated with irreversible LSD1 inhibitor DDP26095 (10 uM), produced by the IEO Drug Discovery Unit–Drug Development Program (IEO DDU-DDP), for 24 h (acute inhibition) or 7 days (chronic inhibition). To induce a constitutive knock-down of LSD1, hiPSC lines were transduced with two different validated pLKO.1 TRC vectors containing shRNAs (sh1 (GCTACATCTTACCTTAGTCAT, Clone Cat. N. TRCN0000046071), sh2 (CCACGAGTCAAACCTTTATTT, Clone Cat. N. TRCN0000046072) against *kdm1a* and maintained with mTeSR-1 medium with puromycin (1 ug/ml) for selection. A pLKO.1 TRC vector containing a scrambled short hairpin sequence was used as a negative control.

### Chromatin accessibility analysis by ATAC-seq

ATAC-seq experiment was performed on fresh fibroblasts in 2 biological replicates per individual according to published protocol^33^. Briefly, 50.000 cells were harvested and washed once in cold PBS for 5 min at 500 g at 4C. The pellet was gently resuspended in 50ul of cold lysis buffer containing 0.1% NP-40 and immediately spun down at 500 g for 10 min at 4 C. Transposition reaction mix containing transposition buffer and 2.5 ul of Tn5 enzyme per sample (Illumina) was added to the pellet and incubated for 30 min at 37 C. The reaction was stopped and DNA was purified using a Qiagen MinElute kit and eluted in 10ul of the EB buffer. DNA was PCR amplified for a total of 11-13 cycles using barcoded primers (Illumina Nextera XT Index Kit v2) and purified using Ampure beads (1.4:1 beads:sample ratio). The quality and concentrations of the eluted libraries was determined using an Agilent Bioanalyzer HS chip. Libraries were sequenced on Illumina NextSeq500 in paired-end mode.

### ATAC-seq data analysis

ATAC-seq data were processed following the steps in^34^ using the custom Snakemake^35^ pipeline. Briefly, reads were trimmed using Trimmomatic^36^ and aligned to the human genome (hg19) using Bowtie2 with parameters -X 2000 --very-sensitive. The read start sites were adjusted due to the transposon insertion specifics of Tn5. Mitochondrial reads and duplicated reads were removed with Picard tools, peak calling was performed using macs2 with parameters –nolambda --nomodel --slocal 10,000.

To look for differentially accessible regions a consensus peak set using samples from all individuals and replicates was produced using the R package DiffBind (minimum of 2 samples to retain a peak), resulting in 96,467 peaks. The accessibility signal was corrected with loess and significantly differentially accessible regions were defined by an FDR threshold of 0.05.

### ChromHMM analysis (for Extended Data Fig.1b)

To analyze the distribution of differential ATAC-seq peaks across different chromatin states we downloaded a 15-state core ChromHMM model (computed on foreskin fibroblast data, cell type E055) https://egg2.wustl.edu/roadmap/web_portal/chr_state_learning.html. The enrichments of peaks (differential peaks more accessible in LSD1^MUT^ or less accessible in LSD1^MUT^ and non-differential peaks) were calculated using the ChromHMM software v.1.22 using the OverlapEnrichment command and default settings http://compbio.mit.edu/ChromHMM/.

### TF activity analysis

Differential TF activity between LSD1^MUT^ and LSD1^WT^ was analysed as in^37^ using diffTF tool^14^. Briefly, this method uses predicted TF binding sites defined by a genome-wide PWM scanner and overlaps these with accessible chromatin in ATAC-seq peaks. As a PWM database source, we used Jaspar CORE 2018 database (Khan et al., 2018), extended TF binding sites by 50 bp in each direction and calculated a fold change in ATAC-seq reads between LSD1^MUT^ and LSD1^WT^ conditions, such that the final TF activity value for each TF corresponds to the mean of genome-wide differences in accessibility at the TF’s binding sites between the conditions. Statistical significance was calculated as a Cohen’s distance of the weighted mean differences distribution to the matching distribution calculated for a permuted binding site of the corresponding TF. Gene ontology analysis on significant TFs was performed using clusterProfiler R package^38^ and all TFs as a background.

**Ageing-related TFs** (as well as other genes in the subsequent RNA-seq analysis of the fibroblast data) were excluded from the fibroblasts analysis. The lists of ageing-related genes were taken from a curated database of human ageing genes GenAge https://genomics.senescence.info/genes/allgenes.php and from a list of genes whose expression was found significantly differential in ageing skin^39^.

### Gene expression profiling by RNA-seq

Samples for all RNA-seq experiments in this work were collected in two biological replicates from each of 2-3 clonal lines or individuals in case of primary fibroblasts (except for the VPA-treated endoderm where individual cell lines were used as biological replicates). RNA was isolated from 600.000-1.200.000 snap-frozen cell pellets using Qiagen RNeasy Mini kit and treated with TURBO DNA-free kit (Ambion) to remove DNA. The quality of RNA was verified on Agilent Bioanalyzer. 1ug of total RNA was used as input for poly-A selection (NEBNext PolyA mRNA magnetic isolation module), cDNA synthesis and library preparation with NEBNext Ultra RNA Library Prep Kit for Illumina according to the manufacturer’s instructions. Library quality was determined using Agilent Bioanalyzer and concentration was measured by Qubit fluorometer. Libraries were sequenced on Illumina NextSeq500 in single-end mode.

RNA-seq reads were adapter trimmed using Trimmomatic and aligned to the human genome (hg19) with STAR aligner version 2.5.2b (https://github.com/alexdobin/STAR) using default parameters. Gene counts were produced with a summarizeOverlaps function from the GenomicAlignments R package and Gencode version 19 annotation of all human genes. Differential expression analysis was performed using an R package DESeq2^40^ in each cell type separately (fibroblasts, hiPSCs, differentiating endoderm or mesoderm or neuronal cells) between LSD1^MUT^ and LSD1^WT^ cell lines. Fibroblasts: samples from primary cell cultures of two individuals grown in two different flasks were used as biological replicates (2×2=4 replicates per condition); iPSCs and VPA-treatment endoderm: 2-3 clonal lines derived from each individual were used as biological replicates; untreated endoderm and mesoderm and neuronal lineages: 2-3 clonal lines derived from each individual were grown in two flasks each and used as biological replicates(LSD1^MUT^: 2 x 2flasks x 2individuals = 8 replicates, LSD1^WT^: 2 x 2flasks x 1individual + 3 x 2flasks x 1individual = 10 replicates). Significantly differential genes were defined by adjusted p-value cutoff < 0.1.

Gene ontology enrichment analyses were done with clusterProfiler R package^38^ using MSigDB database of Reactome-Kegg pathways or “Biological process” GO annotation, p-adjusted cutoff < 0.1.

### Integration of RNA-seq data of LSD1 inhibition and knock-down of normal hiPSCs

Differential gene expression analysis on all hiPSC samples (from the RNA-seq experiments on patient-derived hiPSCs, as well as on normal hiPSC lines treated with LSD1 inhibitor or sh-RNA against LSD1) was performed using edgeR package^41^ and glmRLT function with a model.matrix(∼experiment+condition2), where “experiment” refers to LSD1^MUT^/LSD1inhibition/LSD1knockdown and “condition2” is control samples vs. any of the perturbations. Significantly differential genes in at least one of the conditions were defined by FDR < 0.05. For clustering, the edgeR log fold changes per condition were then sqrt-transformed (i.e. ‘sign(x)*sqrt(abs(x))’) to reduce the impact of a few very large fold changes, and clustered using cluster::clara with k=8. Two of the clusters were merged based on visual similarity and one cluster was not included in the visualization due to the very low number of genes it contained (only two genes), which resulted in 6 final clusters on Fig 4b.

### Integration of publicly available H3K27ac ChIP-seq data

To estimate the levels of histone acetylation at the iPSC state across different gene clusters from Fig. 4b we downloaded raw H3K27ac ChIP-seq data of a matching cell type (human embryonic stem cells, cell line H1) from the Roadmap Epigenomics Project https://www.ncbi.nlm.nih.gov/sra/SRX012366[accn]. The raw sequences were adapter trimmed (using Trimmomatic v.0.33^36^) and aligned to hg19 using bowtie2 (v.2.3.2^42^) with parameters -X 2000 -t. We then calculated the levels of H3K27ac at the enhancers linked to the genes in six clusters from Fig. 4b using the DiffBind R package (version 3.0.15^43^), which we visualized as density plots in Fig. 4c.

### Gene regulatory network analysis

Gene regulatory network was constructed from the matched RNA-seq and ATAC-seq data of healthy human samples and cell lines described in Table 1 using an R package GRaNIE https://doi.org/doi:10.18129/B9.bioc.GRaNIE. The studies were selected to include relevant cell types for the CPRF syndrome (brain tissues, differentiation time course of pluripotent stem cells, mesoderm lineage) and have paired RNA-seq and ATAC-seq data from the same samples of normal cells or tissues (unmutated/unperturbed). Raw input data was downloaded from the original publications and uniformly processed with the same pipelines for RNA-seq and ATAC-seq processing, as the fibroblast data described above. The resulting consensus set of ATAC-seq peaks across all 63 samples (peaks retained in the final peakset if they were identified in at least 2 samples) includes 500,000 peaks and counts within this consensus peakset were used as input for the network construction.

The computational approach to construct a gene regulatory network consists of two steps: building TF-enhancer links and enhancer-gene links. The details of this approach are described in^15^. Briefly, in the first step the expression of each TF was correlated with accessibility of each of the enhancer regions (=ATAC-seq peak) with and without a known binding site of the TF (foreground and background, respectively). Known binding sites were defined using the HOCOMOCO database v.10. Significantly correlated TF-enhancer links were identified using empirical FDR of 20% (calculated separately for each TF) and an absolute correlation Pearson’s coefficient > 0.4. In the second step chromatin accessibility at the ATAC-seq peaks was correlated with the expression of all genes < 200kb away from the peak and peak-gene links were retained if they were positively and significantly (P < 0.05) correlated (our assumption is that accessibility at the regulatory region positively correlates with expression of the linked gene), and if their Pearson’s correlation coefficient was > 0.4. This approach resulted in a total of 1.3 million TF-enhancer-gene links, connecting 393 TFs with 15,128 unique genes via 96,050 unique ATAC-seq peaks.

### TF regulons expression analysis in LSD1^MUT^ vs. LSD1^WT^ cells

TF regulons were defined as all TF-gene links of each TF within the network. To identify which TF regulons were affected by LSD1^MUT^ we calculated aggregated mean expression log2 fold change of each TF regulon in LSD1^MUT^ vs. LSD1^WT^ and tested whether it differs significantly from all expressed genes (background) in the corresponding cell type (two-sided Student’s t-test). The raw p-values from the test were corrected for multiple testing using Benjamini-Hochberg approach and TFs which passed the FDR < 0.1 were defined as having significantly affected regulons (Fig. 2c and 3d, Extended Data Fig. 2a). The difference in means between TF regulons and background genes’ expression was further used in the scatter/density plots and in the heatmap (Fig. 2d and 3d-e).

In the VPA treatment experiment significantly affected TF regulons were defined in untreated endoderm and their expression was calculated in VPA-treated samples, preserving the classification into significant and other TFs to keep the comparison between the same sets of TFs before and after treatment.

### LDSC analysis

We performed stratified LDSC (S-LDSC^20^) to identify if any of the TF clusters were enriched for heritability in GWAS traits. GWAS summary statistics for 806 traits that included participants of European descent were downloaded from the GWAS catalog (https://www.ebi.ac.uk/gwas/) and harmonized. The summary statistics were converted into the LDSC format as described on the LDSC github repository (https://github.com/bulik/ldsc). Summary statistics with fewer than 10,000 individuals or less than 100,000 genetic markers were removed, leaving 442 GWAS traits. We downloaded 54 sets of genomic features from the Alkes Group public repository (https://alkesgroup.broadinstitute.org/LDSCORE/) to serve as a baseline for the enrichment analyses as described (22). Peak sets were created by taking the enhancer regions of the full LSD1 GRN, as well as the enhancer regions connected to the TFs of clusters 1-3 and those connected to the top 10 TFs for the neuronal, endoderm and mesoderm lineages. The peak set for the full LSD1 GRN was added to the background regions, while the other enhancer regions were tested for enrichment of heritability. The LDSC enrichment P-values were corrected for multiple testing within each trait using the Benjamini Hochberg method.

### Immunoblotting

Cell lysates were separated on precast 4–12% Bis-Tris SDS-PAGE gels (Thermo) in MES buffer and transferred onto nitrocellulose membranes. The blots were incubated with primary antibodies overnight at 4° C and for 1 hour at RT with HRP-conjugated secondary antibody. Membranes were developed using the Immobilion Western Chemiluminescence HRP substrate (Millipore). In this work we used the following antibodies: LSD1 - Santa Cruz, # sc-271720;

H3K27ac - Active Motif, #39133; H4 - abcam, # ab10158.

## Acknowledgments

We thank the staff at the EMBL Sequencing Core Facility for sample preparations and data generation. We thank Nichole Diaz for reprogramming fibroblasts and differentiating hiPSCs and Guy Riddihough for the critical feedback on this manuscript. D.B. was supported by a fellowship from the EMBL Interdisciplinary Postdoc (EIPOD) programme under Marie Sklodowska-Curie Actions COFUND (grant agreement number 664726). This work is supported by the DFG fund (SPP 1738 to K.M.N.).

## Author contributions

D.B., J.B.Z and K.-M.N. conceived the study. J.B.Z and K.-M.N. jointly supervised the study. D.B. performed ATAC-seq and RNA-seq experiments under the supervision of K.-M.N. P.-L.G., A.L.T. and A.Ad. performed LSD1 inhibition and knock-down experiments in healthy hiPSCs, generated and analysed resulting RNA-seq data under the supervision of G.T. N.F.-N.M. helped with cell culture and performed Western Blotting experiments under the supervision of K.-M.N. A.Ah. performed gene expression analysis on LSD1 differentiations and processed input data for the eGRN construction under the supervision of D.B. and J.B.Z. C.A. helped with eGRN analysis, A.C. and M.C.L. performed LDSC analysis under the supervision of J.B.Z. D.B. conducted all other pre- and post-processing computational analyses, data integrations and visualizations under the supervision of J.B.Z. and K.-M.N. S.R. and V.N. obtained primary patient samples with the help of P.L. and K.M.P. D.B., K-M.N. and J.B.Z wrote the manuscript, with the input and approval of all co-authors.

