## Supplementary Figures 1-4. for "Pathological LSD1 mutations repress enhancer-mediated gene regulatory networks in early differentiation"

Supplementary material

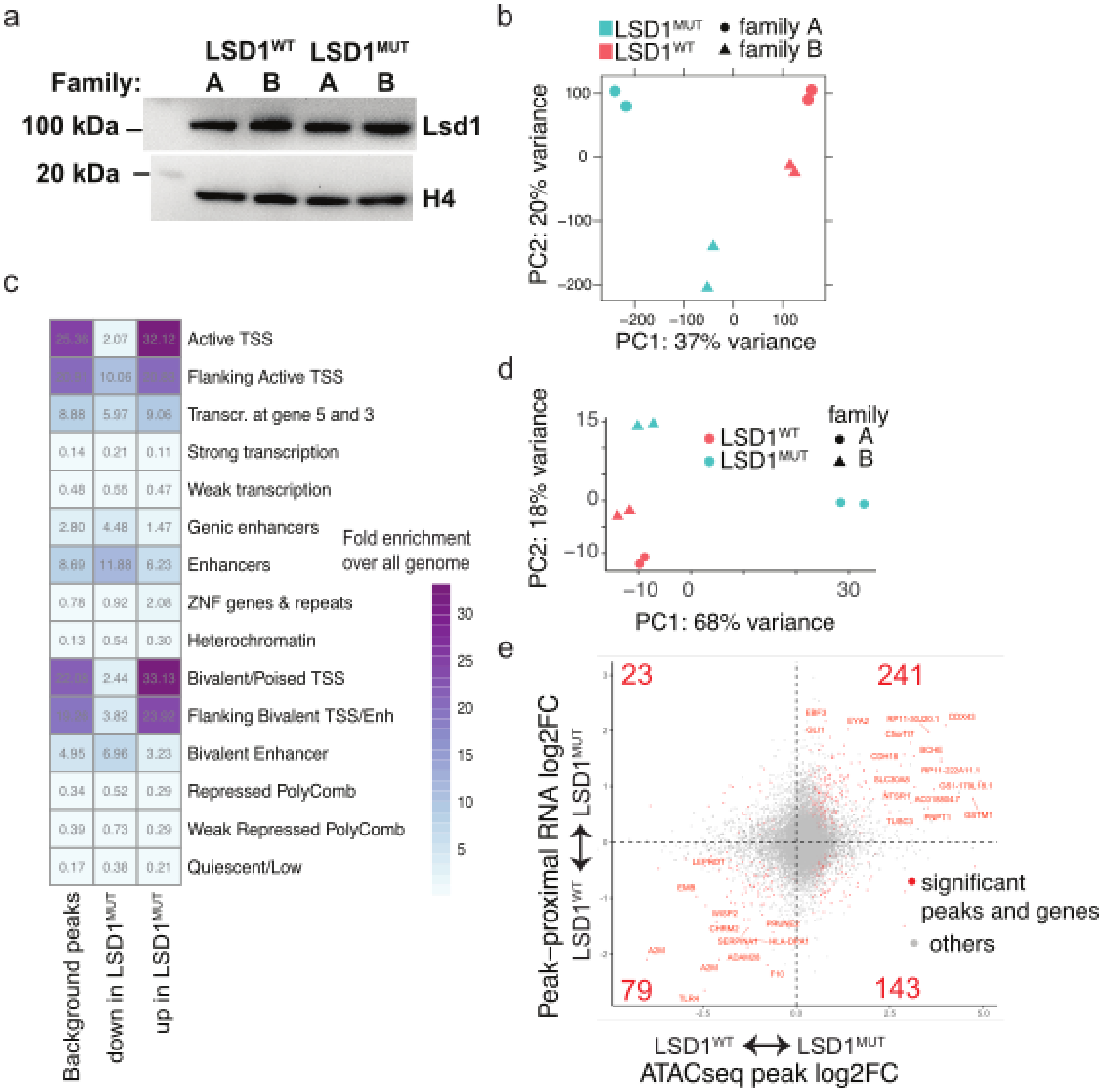


***Extended Data Fig.1.*** *Effects of LSD1 mutation on chromatin accessibility and gene expression in fibroblasts. a) Western blot of nuclear extracts from primary fibroblasts showing Lsd1 protein expression (92 kDa size) across samples and replicates. b) Principal component analysis (PCA) plot of ATAC-seq peak accessibility data in primary fibroblasts. Principal components 1 and 2 are shown together with the variance they explain. c) Enrichments of non-differential ATAC-seq peaks and significantly up- or down-regulated peaks in LSD1$^{MUT}$ in 15 chromatin states identified in the Roadmap epigenome project for foreskin fibroblasts (E055). Fold enrichments relative to the whole genome background were calculated with ChromHMM software [http://compbio.mit.edu/ChromHMM/](http://compbio.mit.edu/ChromHMM/). d) PCA plot of RNA-seq data in patient-derived fibroblasts displaying first two principal components. e) Scatter plot showing correlations between ATAC-seq peak accessibility and gene expression changes in LSD1$^{MUT}$, where peaks were linked to the nearest gene within 3 kb from the transcription start site. Significantly differentially accessible peaks and differentially expressed proximal genes are shown in red.*

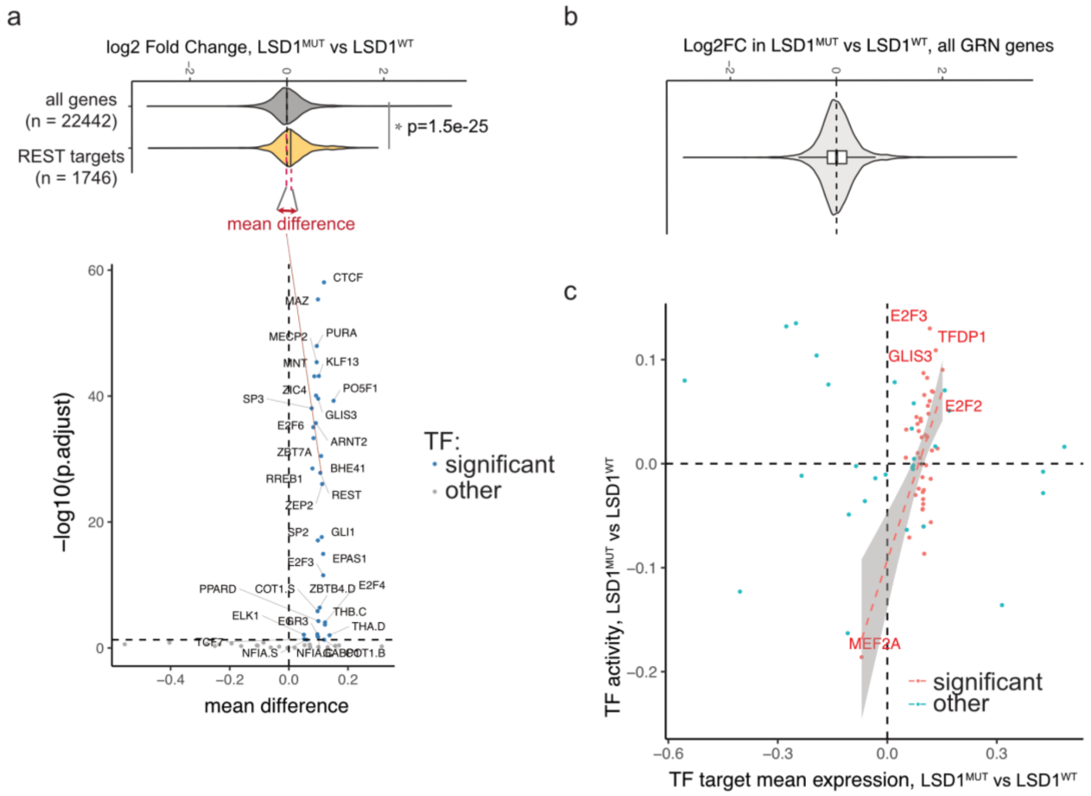


***Extended Data Fig.2.*** *a) Violin plot showing that expression of the REST regulon (its target genes) is on average increased compared to all expressed genes in fibroblasts and the difference is calculated as mean(REST regulon) - mean(all genes). Mean difference for each TF regulon is shown below as volcano plot against adjusted p-value. b) Distribution of gene expression of all unique genes in the GRN in fibroblasts ($LSD1^{MUT}$ vs $LSD1^{WT}$). c) Scatter plot showing correlation of TF activity (defined by diffTF) and TF regulon expression (defined in panel A).*

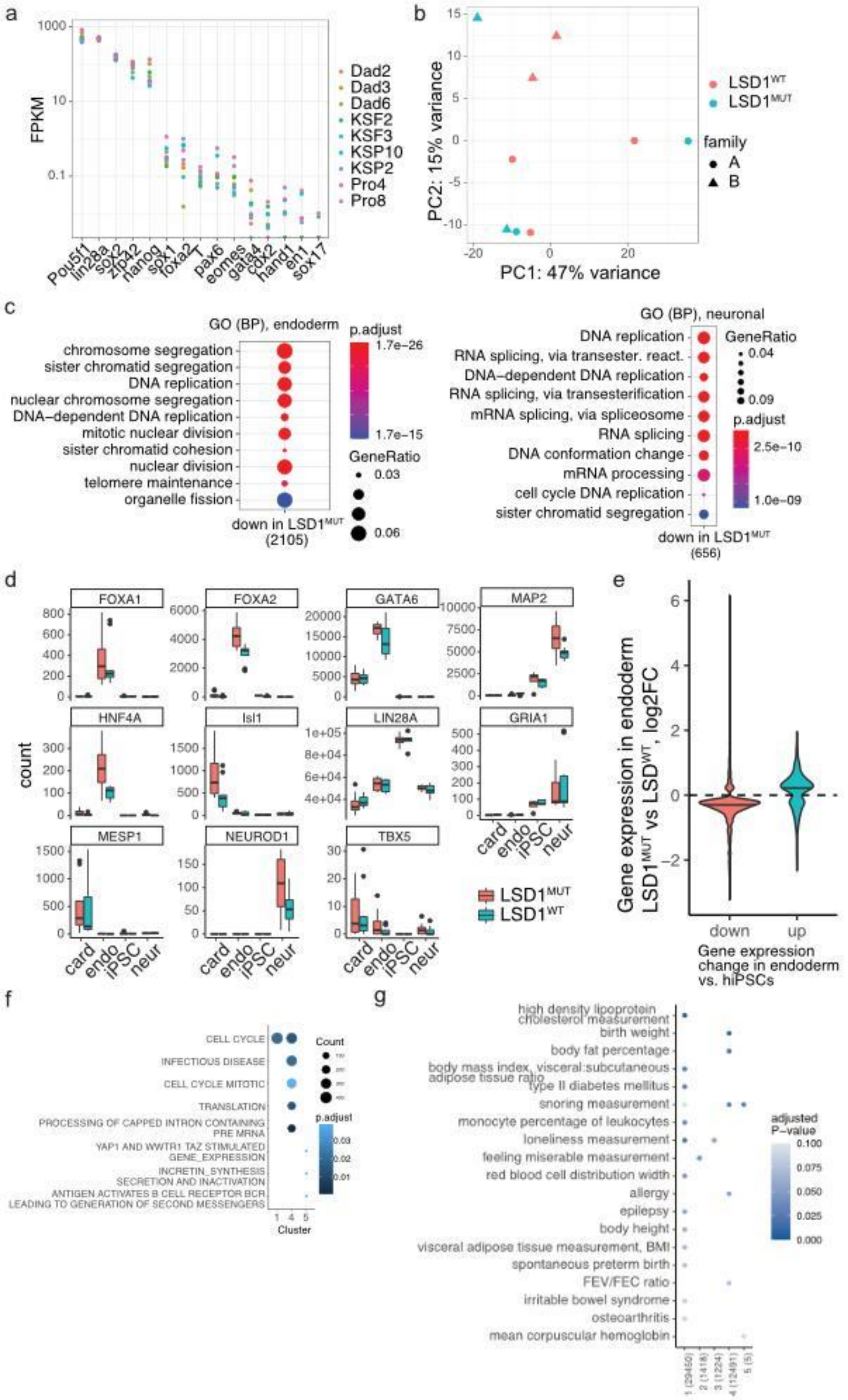


***Extended Data Fig.3.*** *a) Expression of pluripotency markers (Pou5f1, Linc28a, Sox2, Nanog) and lineage markers (Sox1, T, Eomes, Foxa2, Pax6) to confirm the successful reprogramming of primary fibroblasts into hiPSCs: induction of pluripotency markers*

*expression and repression of lineage genes. Expression values are shown as FPKMs from the RNA-seq data. Cell lines: LSD1$^{WT}$ - "Dad" (individual from family 1) and "KSF" (individual from family 2), 2-3 clonal lines per individual; LSD1$^{MUT}$ - "Pro" and "KSP", 2 clonal lines per individual. b) PCA plot of top 300 variable genes in hiPSC lines derived from LSD1$^{MUT}$ and LSD1$^{WT}$ patient samples. c) GO enrichment of significantly differential genes in LSD1$^{MUT}$ lineages, only gene sets with enrichments having adjusted p-value < 0.1 are shown (none for mesoderm lineage). d) Expression of endoderm, mesoderm and neuronal markers in iPSC-derived early differentiated lineages. e) Expression changes of significantly differential genes in LSD1$^{MUT}$ or control endoderm cells, split into two categories: genes up- or down-regulated (by log2FC) in endoderm relative to hiPSCs. The genes are stronger downregulated in LSD1$^{MUT}$ endoderm differentiation compared to control samples. f) gene ontology (GO) enrichment of TF regulons in the three clusters from Fig. 3e. g) LDSC analysis results showing significant enrichments of GWAS traits in regulatory regions associated with TF regulons in clusters 1-5, see Fig. 2e. Number of regulatory regions is indicated in brackets. Adjusted p-values were calculated within each trait while including all GRN peaks as background.*

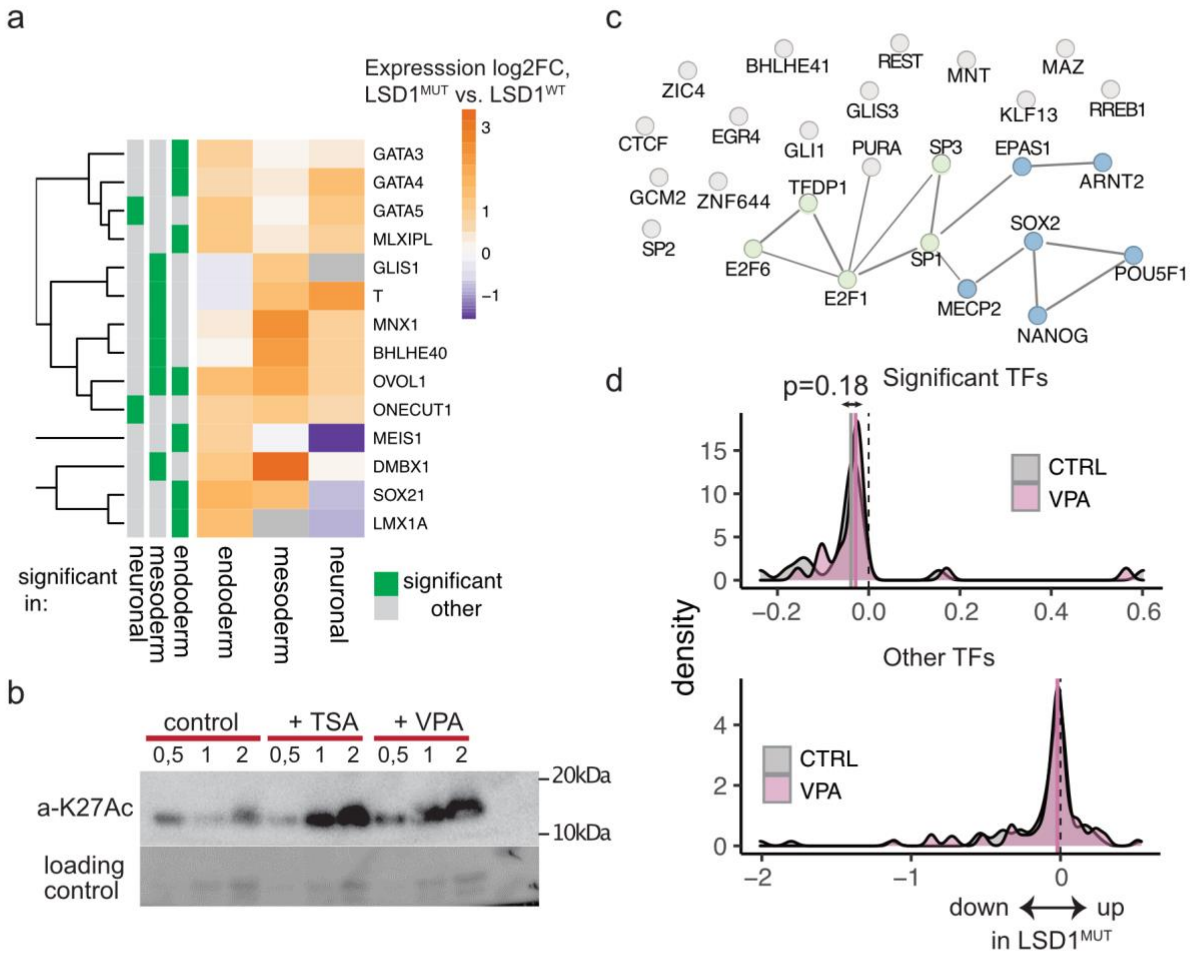


**Extended Data Fig. 4.** a) TF own expression in differentiation, log2 fold change between LSD1$^{MUT}$ and LSD1$^{WT}$. Significant expression changes are indicated in green on the left columns. b) Western blot showing increased H3K27ac levels in patient-derived hiPSCs treated with HDAC inhibitors TSA and VPA (400 uM). c) Protein-protein interaction network of the top 30 TFs linked to the VPA-rescued genes (STRING database), edges correspond to experimental and database evidences. k-means clustering identified 2 groups of TFs shown in colors. d) Distributions of TF regulons expression changes in $LSD1^{MUT}$ vs. $LSD1^{WT}$ in control or VPA-treated endoderm (grey or purple color, respectively). TFs are split into those whose regulons are significantly affected in expression (top plot) or others (bottom plot). Average expression change across all TF regulons in control or VPA-treated endoderm are indicated as dashed vertical lines.
